# London taxi drivers leverage regional boundaries to optimise route choices and improve their navigation skill across three decades

**DOI:** 10.1101/2024.10.31.620595

**Authors:** Christoffer J. Gahnstrom, Sarah C. Goodroe, Stephanie de Silva, Eva-Maria Griesbauer, Jeremy Morley, Ed Manley, Hugo J. Spiers

## Abstract

The world is defined by boundaries. They segment our experience of time and space, and in virtual environments have been shown to impact navigational choices. Here, we test the impact of boundaries on route choices in a real-world environment (London, UK) with a group of expert navigators: licensed London taxi drivers who are required to memorise the layout of over 26,000 streets to obtain their licence. After presenting photographs of a start location and a goal location, taxi drivers were asked to either accept or reject a third target street as forming part of the direct route or not. Performance increased across the adult life-span period in this group (age: 34 to 67). Taxi drivers were faster and more accurate when the target location formed part of a street network boundary (e.g. streets on the edge of the London neighbourhood Soho). Our results are consistent with taxi drivers exploiting the graph structure of the street network to plan routes, as well as consistent with the formation of hierarchical state representations to reduce the dimensionality of the planning problem. Taken together, we show that navigational skill can improve over decades of exposure and that experts exploit regional boundaries for optimal choices, providing a scaffolding over which to form action plans.

## Introduction

The mechanisms that mediate initial acquisition of expert-level knowledge are thought to be necessary for further improvement (Ericsson & Lehmann, 1996). Through years of study, and use of a range of mnemonic devices, prospective London taxi drivers learn to flexibly enact on a remarkable volume of spatial information (Griesbauer et al., 2021). Moreover, the acquisition of the knowledge of the London street network has direct consequences for brain adaptation. In a study by Woollett and Maguire (2011), only the trainee taxi drivers who successfully passed the ‘knowledge exam’ to become a licensed London taxi driver demonstrated increased grey matter volume in bilateral posterior hippocampus. However, it is still unknown if the spatial memory abilities of London taxi drivers continue to improve as they gain more experience. And do they exploit environmental features such as city street boundaries to accomplish these improvements?

Boundaries in spatial environments are known to cause distortions of distance and direction estimations (Hartley et al., 2004; Brunec et al., 2017; Brunec et al., 2018; Stevens & Coupe, 1978; Arnold et al., 2016; Bellmund et al., 2020; Marchette et al., 2017) as well as impairments in memory recall (Horner et al., 2016; Buckley et al., 2022; Radvansky & Copeland, 2006). Despite this, people have the remarkable capability of flexibly navigating through large real-world environments. This ability is thought to be supported by spatial memory systems in the medial temporal lobe and cortex (Maguire et al., 2006; Spiers & Maguire, 2006; Epstein et al., 2017; Bicanski & Burgess, 2020) and to steadily decline with healthy ageing (van der Ham & Claessen, 2020). Certain disease states such as Alzheimer’s dementia are especially linked to rapid decline in real-world navigation ability (Coughlan et al., 2018). Other research also suggests boundaries may enhance memory capabilities by segmenting experienced events (Pettijohn et al., 2016; Barnett et al., 2024). Theoretical accounts of large-scale planning suggest boundaries may be used to segment a road network into discrete regions of space, facilitating a hierarchical representation which drastically reduces the computational demand of planning (McNamee et al., 2016; Earle et al., 2017). The usefulness of boundaries may only become apparent when the environment grows to a sufficient size. However, it remains unsolved whether boundaries are beneficial or detrimental to spatial navigation performance in large real-world environments.

The medial temporal lobe, a brain region known for its important role in spatial and episodic memory, also exhibits neurons encoding distance and direction of boundaries (Lever et al., 2009; Stewart et al., 2014). Boundaries also disrupt the spatial location code of environments in rats and humans (Krupic et al., 2015; He & Brown, 2019). The usefulness of boundaries can be appreciated in large state-spaces where they may form a scaffolding for decision-making and planning. For instance, efficient image algorithms solve a generated jigsaw puzzle by utilising the most salient boundaries - corner pieces (Nielsen et al., 2008). We do not understand how humans leverage boundaries in real-world environments. In particular, how might boundaries in a city impact route choices? And how might those route choices improve with experience, not within months of training, but rather over decades?

The environmental size tested in laboratory settings are usually limited by the amount of learning feasible within the scope of one to a few experimental sessions. By utilising a group of participants that have spent years rigorously learning the same large spatial environment, we can uncover necessary representations to accomplish large-scale planning. Recent work has begun to unravel the principles that may underlie human decision-making and planning when traversing large state-spaces (van Opheusden et al., 2023; Wu et al., 2018; Ho et al., 2022) but these are still far removed from the size of real-world state-spaces such as finding an efficient path between a start and goal location in an environment with millions of possible routes. Fortunately, this is a daily problem faced by licensed London taxi drivers (Woollett et al., 2009) who spend on average 3-4 years studying the London street network to pass their qualifying exam (Griesbauer et al., 2022a).

London taxi drivers have formed an important part of spatial memory research with several landmark studies suggesting the brain adapts to planning demands in large state-spaces (Maguire et al., 2000; Woollett & Maguire, 2011; Woollett, Spiers & Maguire, 2009; Maguire, Woollett & Spiers, 2006) and engagement of brain regions during navigation of a London (Spiers and Maguire, 2006a,b, 2007). One study identified posterior grey matter volume of hippocampus of qualified London taxi drivers was larger than age/gender matched non-taxi drivers (Maguire et al., 2000). They also found that years of experience operating as taxi drivers increased the volume in the right posterior hippocampus. Maguire et al. (2006) replicated the initial finding that posterior grey matter volume positively correlated with years of experience, and also found the anterior hippocampus was negatively correlated with years of experience. By comparing taxi drivers to bus drivers they found that, despite both groups spending time driving in London (year of this matched across groups), taxi drivers had large posterior hippocampus grey matter and small anterior grey matter compared to bus drivers. Woollett, Spiers and Maguire (2009) found that retired taxi drivers showed a slightly reduced posterior hippocampal volume compared to full time drivers, consistent with posterior hippocampal volume being linked to driving a taxi and spatial planning demands. Taken together these studies provide evidence for a link between hippocampal grey matter volume and experience as a licensed London taxi driver. However, in the general non-expert population, spatial ability and brain volume correlations are less consistent further motivating the need to understand the special case of London taxi drivers (Weisberg & Ekstrom, 2021; Brunec et al., 2019; Chrastil et al., 2017; Weisberg et al., 2019; Clark et al., 2020). It is still unknown if these structural brain changes in experts also result in different spatial representation strategies such as forming hierarchical state segmentations and utilising neighbourhood boundaries for planning. Given the grey matter increase over years of experience, our prediction is that the benefit of these putative spatial strategies also scale with experience.

The initial challenge is to quantify what aspects of the street network taxi drivers consider to form boundaries. Griesbauer and colleagues (2022b) asked London taxi drivers to draw areas considered to form part of regions or neighbourhoods on a map of central London. They identified a series of consistently labelled streets as boundaries. Some of these boundaries formed part of well-known neighbourhoods, for instance the streets surrounding Soho in London. In a follow-up study, a different set of taxi drivers were given the instruction to plan routes from a list of start and goal locations and to call out the street names one-by-one that formed part of the participants’ planned route (Griesbauer et al., 2024). Streets which formed part of a boundary had reduced response times suggesting this separate group of participants utilised the same set of boundaries in a process of hierarchical route planning (Fernandez Velaso, Griesbauer et al., 2024). However, there were some limitations such as heterogeneity of street names chosen on routes across participants and small number of routes given the time constraints. Several conceptual approaches to boundary identification have been proposed based on street network segmentation, constructed using taxi trajectories (Manley, 2014) and street network geometry (Filomena, et al., 2019).

To further understand the role of boundaries in large environments, and to address previous limitations, we designed a novel experiment to test if spatial route choice accuracy of London taxi drivers is improved or impaired by boundaries in the street network. This paradigm maximised the number of routes tested and the sampling of a set of variables previously found to affect response times of route planning. These included number of turns, route distance, major roads, and routes crossing neighbourhood boundaries (Griesbauer et al., 2022).Boundaries surrounding these regions can also be considered bottlenecks in the road network where traffic is restricted to a relatively smaller number of alternative roads (McNamee, 2016). For example, the river Thames creates a clear boundary in the street network where each bridge can be considered a bottleneck. To test this we created a route planning task that required deciding whether a probed location was on or off the route between a given start and end location. Start, end and probe locations were selected to allow us to examine the impact of a variety of spatial and street network conditions (e.g. on, off, across boundary) on route choice and reaction time.

We hypothesised that regional boundaries would improve response times and accuracy of accepting or rejecting a street laying on the direct route between two locations in central London. We further hypothesised that experience, and not age, would improve accuracy in our spatial memory and planning task. This study aims to improve our understanding of the remarkable human ability of large-scale route planning in ecological settings.

## Methods

### Task

We designed a novel route planning task using 450 distinct locations scattered around London. Google street view images were extracted from Google Earth (high-resolution 4800×2643 pixels) at specific longitudes and latitudes for each location. Images were compressed to reduce loading delays during stimulus presentation, average compression from 6MB to 300KB while maintaining the same resolution. The 450 locations consisted of 150 starting locations, 150 goal locations, and 150 target locations. Images were presented using a custom Python script and the Psychopy software package (Pierce, 2007).

The task was divided up into two components (Figure 1). (1) Drivers were asked to mentally plan the route between a start and goal location just as they would follow the direct line during a knowledge exam (Griesbauer, 2022). We displayed the street view image and name of the start location at the bottom of the screen for 4-8 seconds, uniformly sampled. After the jittered delay, the street view image of the start location was replaced with the street view image of the goal location, while the text of the start location remained at the bottom of the screen. Along with the appearance of the image for the goal location, a text appeared at the top of the screen stating the location of the image. The street view image of the goal location was present on screen for 4-8 seconds, uniformly sampled. (2) We introduced an accept-reject task where each participant evaluated whether a third street, the target street, formed part of the planned route from start to goal. After the jittered delay for the goal location, the image was replaced by a street view image of a target location along with text in the upper third of the screen, showing the name of the street. The target location image was present until a button press response (yes or no) was made using a button box placed by the right hand of the participant. The yes or no response was mapped to either a response made with the thumb or the index finger, counterbalanced across participants. Each trial consisted of both components – mental route planning and accept-reject of target street – and was separated with an ITI of 3.5-4.5s. The trial order of each route was maintained for all participants.

**Figure 1.**
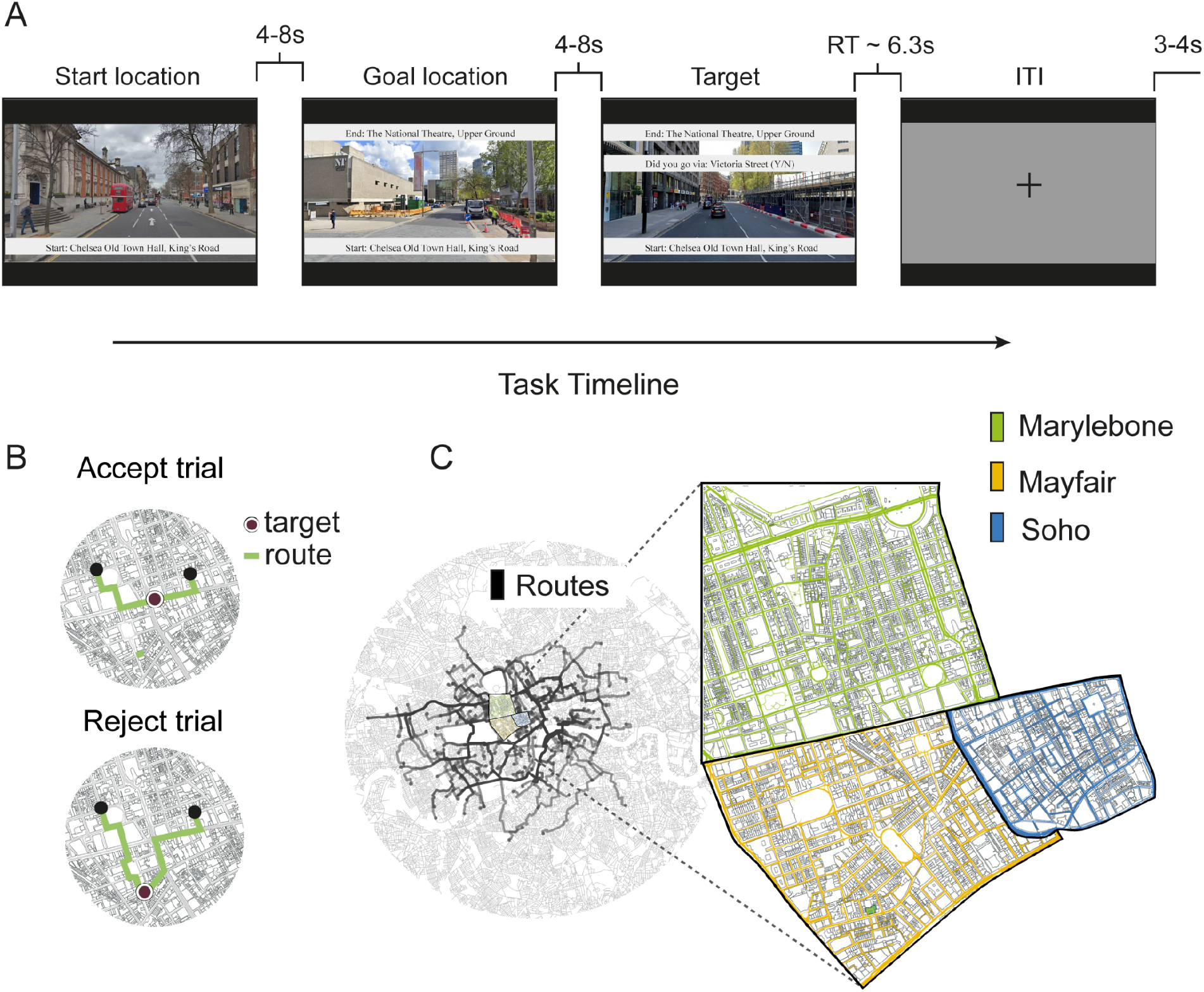
Route choice task. A) Task timeline. Participants viewed a sequence of three street view images taken from unique locations around London (extracted from Google Earth). The task required participants to either accept or reject a target street as belonging to the direct optimal path between start and goal location. The first image is the starting location, and the second image is the goal location. The final image is the target street and the participants needed to make a button response (YES/NO) if this street forms part of the direct route between the first and second image. B) Two example trials where the target street either forms part of the direct route (top; accept trial) or it does not form part of the direct route (bottom; reject trial). C) A map of the street network of london. Each black line represents one of the 120 routes tested. Expanded regions show three example neighbourhoods (Marylebone, Mayfair, and Soho) in the street network with black lines forming boundaries between each region.

Routes were selected which sampled the London street network based on a number of route characteristics. These included the following: route path distance, number of turns, cardinal direction of the route, alignment of route to main axis of street network, target distance to start location, target distance to goal location, detour distance of target, target being on route, target being off route, target being on an A-road (major roads in United Kingdom), and routes crossing the river. The different routes and targets also uniformly sampled the geographical area of the 3-mile radius around Charing Cross, the centre point for the Knowledge (Figure 1).

### Participants

The study was approved by the ethics committee for UCL Division of Psychology and Language Sciences (fMRI/2021/001). For online testing, ethics approval was received from UCL Department of Experimental Psychology (EP/2018/008). Twenty-six participants (age ranging from 34 to 67 years, mean age: 52.1, std age: 9.16, all male, 25 right-handed) completed the full experiment including one training session. All participants were London licensed taxi drivers holding green badges authorised by Transport for London. The average years of taxi driver experience was 14.07 years (min = 1, max = 38, std = 10.35). The participants were recruited using a variety of methods including online advertising in magazines, social media, and radio interviews (BBC radio 4).

### Online training and Questionnaire

Each participant completed 30 training trials at home in the week prior to coming in for the experimental test session. We wanted to familiarise the participants with the mental route planning and the accept-reject aspect of the task. Participants also completed a demographic questionnaire, and a taxi-driver-specific questionnaire regarding their experience and preferences as a driver.

### Experimental Design and Behavioural analysis

Each trial contained an accept-or-reject response for the question “Did you go via: X street?”, where X street refers to a street that may or may not lay on the direct route between a given start and goal location. We collected response times for 120 route planning trials during the experimental test session concurrently with functional magnetic resonance imaging (brain data not presented here). Subsequently, we investigated the consistency of responses across participants and factors that may influence the response times.

We calculated optimal distance paths between each start and goal location pair by first extracting the street network graph of Greater London, UK, from OpenStreetMaps using custom Python code and the osmnx package (Boeing, 2017; Haklay & Weber, 2008). The street network graph contained 568,473 nodes and 1,085,199 edges. Optimal paths were also calculated from start, to target, to goal location. The detour deviation was computed as the fractional increase in distance by going through the additional target street location, and was used in an ordinary least squares, linear regression model to account for response times. A correct accept response was defined as any trial where the detour deviation caused by the target street was less than 10% of the total route length.

We used linear and logistic regression to test our hypotheses of how boundaries in the street network influences accuracy and response times. All response times were log-transformed separately for each participant. The regression was run separately for each participant and the resulting coefficients were used to run a two-sided t-test, testing for factors significantly different from 0.

Our study tested the following two main hypotheses:

1. Target streets placed on regional boundaries lead to improved task route planning task accuracy and faster response times.
2. Years of experience as a licensed taxi driver in London lead to improved route planning task accuracy.

All tested regressors and associated predictions on task accuracy and response times for the first main hypothesis are listed in the table below:

**Table 1.** Experimental task variables. Values for each regressor were calculated for each of the 120 trials. The regressors either applied specifically to a target street characteristic (e.g. target on boundary, target off boundary, and target on A road) or to a route characteristic (e.g. route crossing boundary, route along boundary, or route crossing river). Each regressor was hypothesised to increase or decrease accuracy and response time. Control variables included other possible influences of boundaries on task performance and response times as well as familiarity with street names captured by the target on A road regressor.

| Regressors | Accuracy | Response time | Control |
| --- | --- | --- | --- |
| Number of turns | Decrease | Slower | No |
| Target on boundary | Increase | Faster | No |
| Target off boundary | Decrease | Slower | No |
| Target on A road | Increase | Faster | Yes |
| Route crossing boundary | Increase | Faster | Yes |
| Route along boundary | Increase | Faster | Yes |
| Route crossing river | Decrease | Slower | Yes |
| Target route increase | - | Slower | No |

For our second main hypothesis, we separately tested the influence of years of experience as a licensed Taxi driver on task accuracy and response times using Pearson correlation. There were two confounding factors which we also controlled for, namely GPS use and level of education. These variables were collected in a questionnaire completed by participants before the start of the route choice task. We performed a linear regression analysis with task performance as our main variable of interest and the control variables as covariates.

## Results

### Overall task aim

We designed the experiment to test hierarchical (boundary-based) and non-hierarchical route characteristics influencing flexible spatial memory behaviour of London taxi drivers. Each trial consisted of three street view images presented in succession: 1) the start location image, 2) the goal location image, 3) the target location image (Figure 1). Participants were asked whether the final street view image, i.e. the target locations, were part of the direct line route going from the start to the goal location. Accept or reject answers were indicated by button response and response times were recorded (Figure 2B, 2C). The experiment consisted of two types of route manipulations; varying the characteristics of the start-goal route (e.g. length, direction, crossing the river Thames) and varying the target position relative to the route (e.g. part of the direct route, or variable distance away from of the direct route).

**Figure 2.**
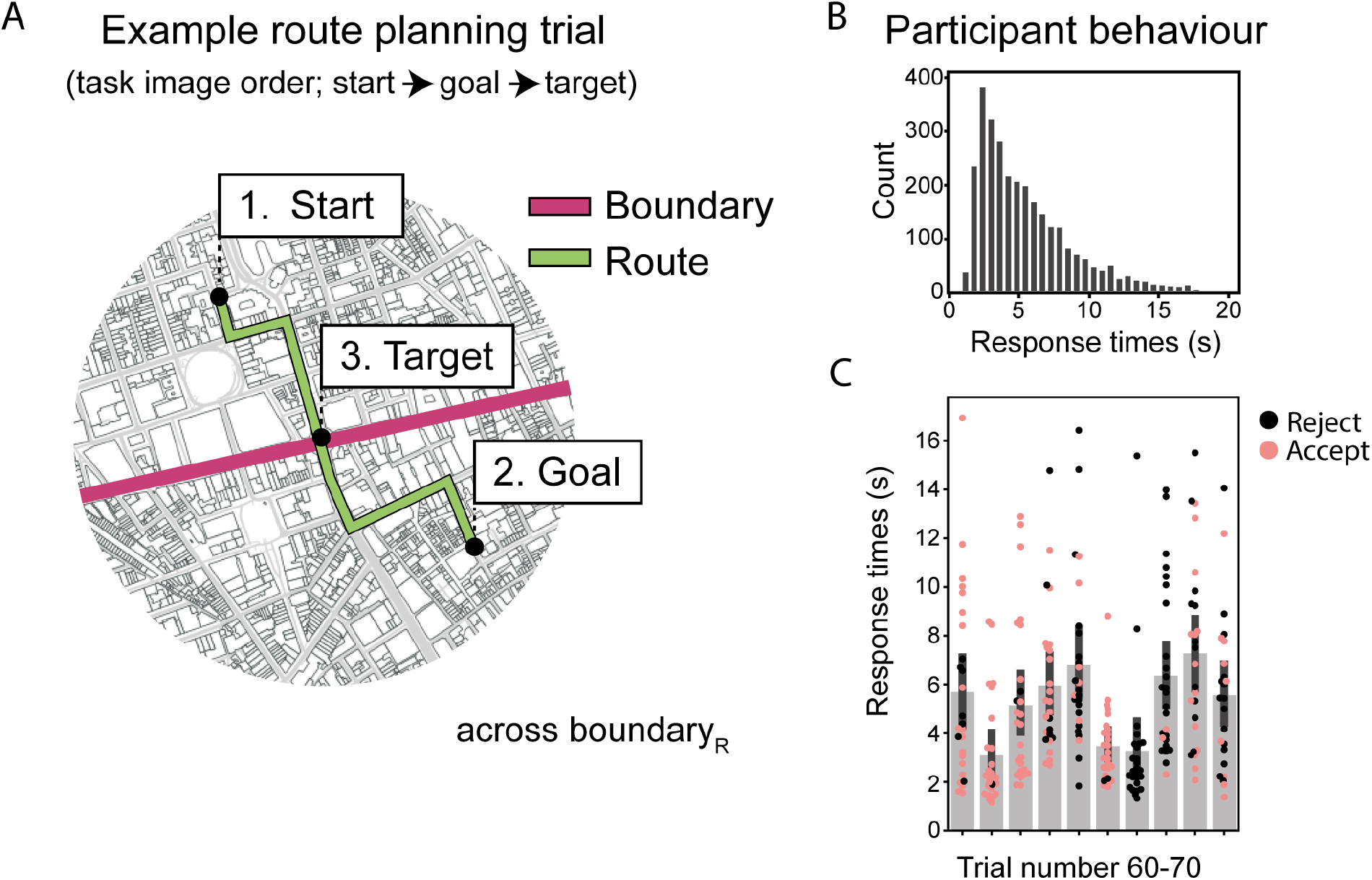
Overview of participant response variability. A: One example route showing the start, goal, and target location overlaid on the London street network. The example route crosses a pre-defined boundary from a separate dataset (Griesbauer et al,. 2021). The target location was selected to be on the direct route between the start and goal locations and on the boundary as defined by the licensed London taxi drivers. B: Response behaviour from all participants in histogram (n=26). C: A subset of trials (60-70) to demonstrate variability across routes and reject or accept choices.

### Taxi drivers utilise regional city boundaries in route choice task

The task was designed to test the utilisation of real-world street hierarchy, in the form of perceived regional boundaries, during route selection. First, we selected all the accept trials, these are trials where the target street increased the route length by a maximum of ten percent to account for slight inaccuracies in computing direct route distances (see methods using osmnx). Together, these contained all trials where target streets lay on the direct route between start locations and goal locations (n=51). Second, we compared the accuracy scores between the trials where the target street was on a boundary with trials where the target was not on a boundary (boundaries as defined in a separate group of taxi drivers from Griesbauer et al., 2021). Participants were significantly more accurate for trials where targets were on a boundary (paired samples t test; t(25)=4.44, p=0.00015; Figure 3).

**Figure 3.**
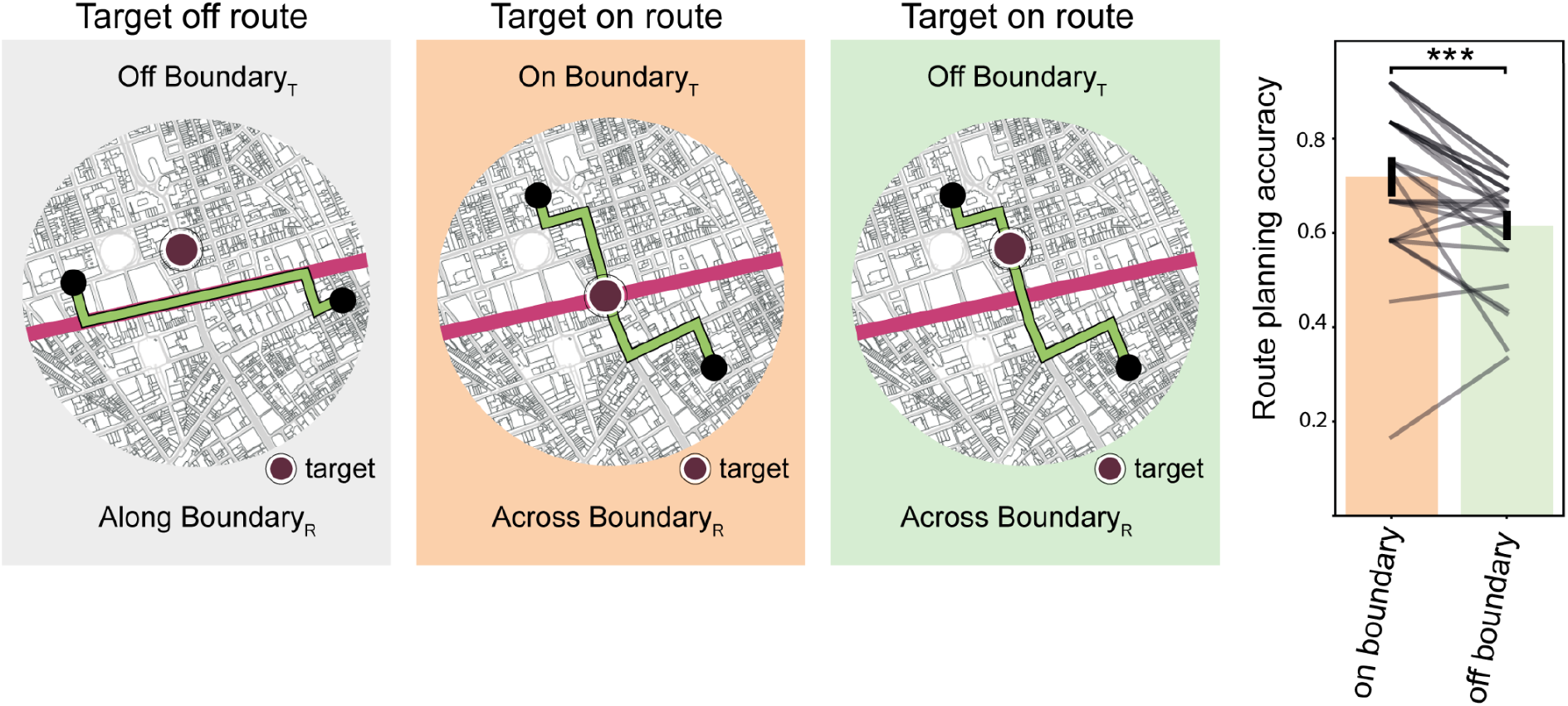
Improved planning accuracy when target streets are on boundaries in the London street network. Left: Schematic for example route (in green) where the target street is off boundary and the route runs along a boundary. Middle: Schematic for an example route (in green) where the target is on boundary and the route crosses a boundary. Right: Schematic for an example route (in green) where the target is off boundary and the route crosses a boundary. Middle and right images show the target as on the shortest route between start and goal locations. Participants are consistently more accurate at the route choice task when the target street is on a boundary, as compared with when a target street is off a boundary (paired samples t test; t(25)=4.44, p=0.00015).

### Boundaries in the street network influence accuracy and response times

A hierarchical representation of the environment is hypothesised to reduce response times and improve choice accuracy when target streets are on boundaries in the street network. Several categorical predictors (number of turns, on boundary, off boundary, across boundary, along boundary, and across river) were first entered into a logistic regression to predict correct and incorrect responses in our route planning task (Figure 4). Second, the same categorical predictors were entered into a linear regression analysis (ordinary-least squares), along with route increase (Figure 4), to predict response times. Each model was fit separately per participant and resulting coefficients were used for population-level tests. Target streets placed on boundaries (bottlenecks in the state-space) significantly improved choice accuracy (t test t(25)=2.11, p=0.045), while target streets not placed on a boundary reduced choice accuracy (t test t(25)=-2.46, p=0.021). Importantly, there was no significant effect on accuracy if a target fell on a major road (A-roads in the United Kingdom), which was considered as a proxy for street familiarity (t test t(25)=-0.23, p=0.82). The number of turns in a route reduced choice accuracy, where more turns resulted in worse performance (t test t(25)=-2.23, p=0.035).

**Figure 4.**
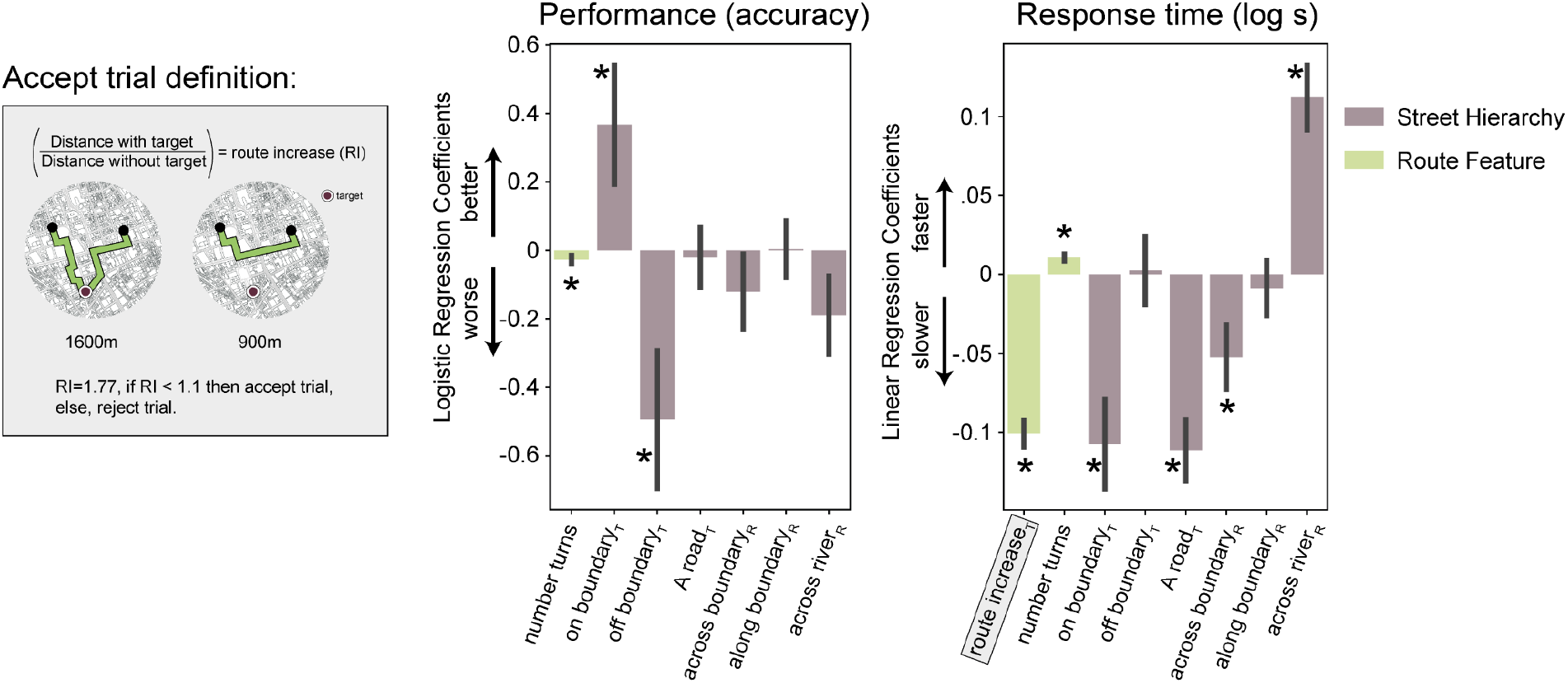
Multiple factors impact accuracy and response times in expert spatial route choices. Accuracy is defined as correctly accepting a target street which falls on the direct route between start and goal location. Target street is allowed to deviate from the direct route by up to 10% in terms of increased route distance. Right: Example single route schematic in grey box where target causes 77% route increase and therefore classified as a reject trial. Middle: Multiple logistic regression identified participants as consistently more accurate at the route choice task when the target street is on a boundary (t test t(25)=2.11, p=0.045), and less accurate when a target street is off a boundary (t test t(25)=-2.46, p=0.021). More turns in a route impaired performance (t test t(25)=-2.23, p=0.035). Right: The influence of street network boundaries and route features on accuracy and response times was quantified using multiple linear regression. Amount of target route increase (RI), target on boundary, target on major road (A-road), and route crossing a boundary, all decreased response times (t test t(25)=-11.95, p<0.00001; t test t(25)=-6.24, p<0.00001; t test t(25)=-5.71, p<0.00001; t test t(25)=-2.59, p=0.016; respectively). The number of turns in a route (t test t(25)=4.79, p=0.000064) and the route crossing the river Thames (t test t(25)=5.48, p=0.000011) increased response times.

**Figure 5.**
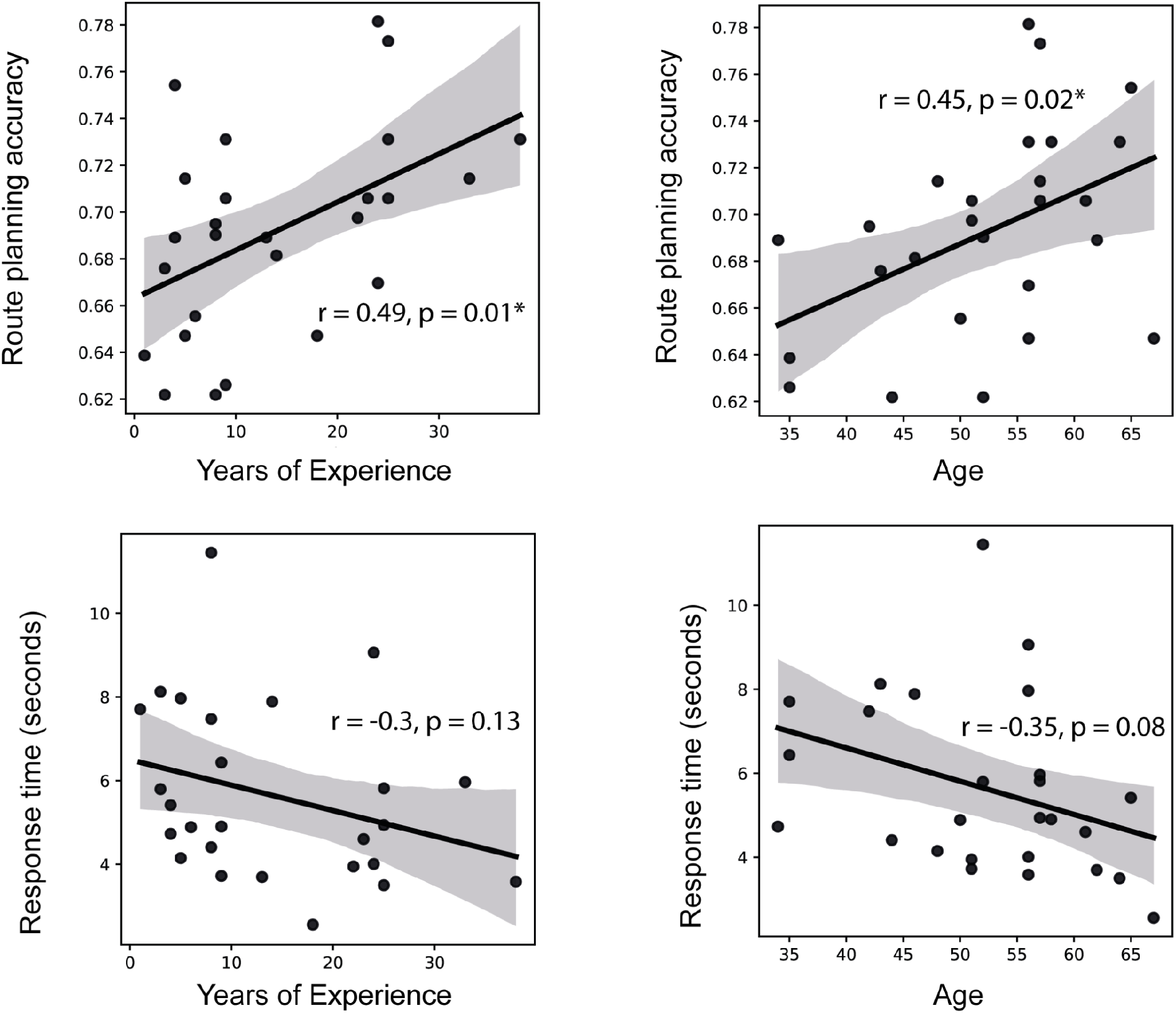
Experience significantly improves choice accuracy and not response times. Top row: Experience as a licensed Taxi Driver correlated with improved accuracy in our task (Pearson’s r=0.49, p=0.01). The age of the participants also correlated with choice accuracy (Pearson’s r=0.45, p=0.02) although given the multicollinearity of age and experience these two factors are easily not separable. Bottom row: Years of experience and age both showed no significant relationship with mean response time in the route choice task (Pearson’s r=-0.3, p=0.13 and Pearson’s r=-0.35, p=0.08, respectively).

Similarly to the impact on accuracy, response times were influenced by boundaries in the London street network. Trials with target streets on boundaries had faster response times (t test t(25)=-6.24, p<0.00001) while target streets off boundaries had no significant impact on response times (t test t(25)=0.12, p=0.91), and when the route crossed regional boundary response times were faster (t test t(25)=-2.59, p=0.016). However, the most significant boundary in the London street network, the river Thames, had the opposite effect and increased response times (t test t(25)=4.48, p=0.00014).

The inclusion of each target street in the route increased the total path length depending on the target street distance from the original route (Figure 4). These increases in route lengths resulted in faster response times (t test t(25)=-12.17, p<5.3e-12), most likely driven by large increase in route distance leading to easy reject trials. The number of turns in a route increased response times (t test t(25)=5.01, p=0.0000014). Our proxy for street familiarity, target streets on major roads, significantly decreased response times (t test t(25)=-5.71, p<0.00001).

### Experience improves accuracy of spatial route choices in expert navigators

Years of experience as a taxi driver correlated with performance in our route choice task (pearson’s r=0.49, p=0.01). We also found a similar relationship for age (pearson’s r=0.45, p=0.02). The regressors for age and experience are collinear, leading to difficulty in separating experience from age (Pearson’s r=0.5, p=0.009). The influence of experience on performance was further considered using linear regression, with years of experience as the main variable of interest predicting performance and controlling for GPS use and level of education. Accuracy in route choices was significantly predicted by years of experience as a licensed London taxi driver (β=0.0023, t(25)=2.81, p=0.01). None of the control variables predicted a significant relationship with performance (GPS use β=-0.0069, t(25)=-0.334, p=0.742; education β=0.0024, t(25)=0.893, p=0.382) and average response times showed no significant relationship with either years of experience or age (Pearson’s r=-0.3, p=0.13 and Pearson’s r=-0.35, p=0.08, respectively).

## Discussion

Humans have the extraordinary ability to form large structured representations, and to use those representations to perform flexible behaviours. Here, we developed a novel route choice task to target the route selection process of expert navigators. The task consisted of presenting a series of three street view images from London in succession, where the final street view image either formed part of a direct route connecting the first two street images, or it did not. Taxi drivers were able to accurately accept or reject the final street view image as forming part of the direct route. Interestingly, this ability positively correlated with years of experience, suggesting route choice accuracy improves over decades paralleling changes in hippocampal grey matter in taxi drivers, as previously reported (Maguire et al., 2000, 2006).

The main hypothesis of the experiment was that taxi drivers utilise boundaries to facilitate the computationally demanding task of route planning. In support of this, taxi drivers had increased accuracy and reduced response times when target streets lay on a boundary compared with when they did not. This result only considered accept trials where the target street lay on a direct route between start and goal location. When we considered all trials in a multiple linear regression, the same finding held. In fact, we found a double dissociation where ‘on boundary’ target streets significantly improved accuracy, and ‘off boundary’ target streets significantly decreased accuracy. One possible confound could be familiarity with street names driving the effect. If boundary streets are more familiar to taxi drivers then perhaps they are more readily available to evaluate during the route selection process leading to improved performance for those streets. As a proxy of street familiarity we included target streets on major roads (e.g. Oxford Street) as a covariate, and found no significant effect on choice accuracy. However, major roads did reduce response times suggesting that they were more familiar but were not used to facilitate route selection in the same way as boundaries. Notably, boundaries and major roads had independent effects on response times.

Previous studies show humans have a propensity to select routes that reduce the number of regional boundaries crossed despite identical route distances (Wiener & Mallet, 2003; LeVinh & Mallet, 2024). In virtual environments, response times are better explained in terms of the number of regional boundaries crossed than the number of steps (Balaguer et al., 2016). Here, we demonstrate the advantage of segmenting street networks into regions in real-world city environments to facilitate the large-scale ecological problem of route planning.

How does this align with previous work showing biases and impairment in performance when human participants segment environments into regions (Burris & Branscombe, 2004; Allen & Kirasic, 1985; Peer & Epstein, 2021)? Although these studies indirectly suggest a cost of segmentation, we suggest that the cost also comes with a benefit. Perhaps the loss of local precision of distance and direction estimates coincides with overrepresentation of boundaries and improved understanding of global regionalisation. Activity in the medial temporal lobe and scene selective brain regions are known to covary with boundary variables (Doeller et al,. 2008; Bird et al., 2010; Lee et al., 2017; Park et al., 2011; Ferrara & Park, 2016; Dilks et al., 2013; Julian et al., 2016; Shine et al,. 2019; Park & Park, 2020; Marchette et al., 2014). One possible interpretation is that spatial navigation ability is built on the backbone of environmental boundaries and humans in particular rely heavily on boundary representations, even beyond reliance on other environmental variables such as distal landmarks (Lee, 2017).

Further work is needed to understand how these regional boundaries may be constructed to begin with. Initial research on London taxi drivers suggests these are based on environmental features such as the rectilinear nature of the designated region (Griesbauer et al., 2022a). The river Thames, which geographically segments London into two split regions North and South, was not consistently identified as a boundary. Our accuracy measurements also support this counter-intuitive finding with no significant change in accuracy if a route crossed the river although it induced a strong increase in response times. Currently it is unclear why most boundaries and major roads lead to faster responses, but bridges over the river Thames lead to slower responses. One possibility is that these locations lead to more consideration of routes compared to other boundaries.

Here we found cross-sectional evidence that over a range of experience from 1 year to 38 years for greater route choice accuracy with experience. In our study age and experience were correlated making it difficult to determine whether the relationship is driven by one or both. Such an improvement with age stands in contrast to many studies that show a reduction in spatial navigation performance with age (e.g. Coutrot et al., 2018; 2022; Lester et al., 2017; Van Der Ham, 2020). Past research with taxi drivers has shown they were better able to construct an accurate representation of an environment shown via film than a matched group of non-taxi drivers (Woollett and Maguire, 2010). But that prior study did not examine the impact of years of experience on performance. Future research linking potential changes in hippocampal size or subfield size with performance accuracy will be useful for exploring whether the advantages observed in performance are linked to changes in hippocampal structure found in previous studies of taxi drivers (Maguire et al., 2000; 2006).

### Limitations and Future Directions

The use of planning in our task is an assumption which may need further consideration. Although unlikely given the performance levels, it is possible that our participants may have employed different strategies or heuristics to solve the task. For instance, they may have used some components of compositionality whereby they recall disjointed memories of navigating through London to roughly similar locations as probed by the start, goal, and target streets (Kurth-Nelson et al., 2023). Our experts were not probed on the complete route sequence of streets as doing so would prohibit the number of trials and sampling of a wide range of hierarchically relevant variables of the street network. A recent study asked a different group of licensed London taxi drivers to call-out each street that lay between a start and goal location. They found evidence in the response times that taxi drivers use predictive representations and hierarchical chunking of transition sequences (Fernandez Velasco et al., 2024). Our approach had the benefit of providing an objective measure of performance while also probing many different instances of boundaries all across London. An intermediary task design could possibly bridge these two approaches by sampling route choices on multiple points along the same route.

We hypothesise that taxi drivers use boundaries and bottlenecks in the state-space to efficiently plan and reduce the computational demands (McNamee et al., 2016). The boundaries were quantified using a separate study where a different set of taxi drivers drew separating lines on a map of London of where they considered these boundaries to be (Griesbauer et al., 2022). In the current study, we consider any given point on these boundaries as a bottleneck in the state-space. However, the boundaries did not have perfect agreement across the tested participants in the original study. It is entirely possible that taxi drivers form partly individualised representations due the certain areas of London they prefer to pick up passengers. Future work can define these boundaries within the same population of taxi drivers to investigate individually mapped hierarchical representations and how they influence spatial memory and planning.

The measurement of accuracy used in our analyses can be challenging to quantify. We made distance calculations using the osmnx software package (Boeing, 2017) which pulls street network information from OpenStreetMap repository maintained by a community of volunteers via open collaboration. This can create some anomalies or inaccuracies given that the street information is based on openly available information. Moreover the street network of London is under constant development with high frequency in road closures. This means that many routes change over time leading to uncertainty for taxi drivers about the current best route between a given start and goal location. Although the influence of this lack of a ground truth on our results cannot be measured, future tools may help in establishing the degree of change in the street network and how taxi drivers update their existing hierarchical representations in the face of new information.

In conclusion, licensed London taxi drivers performed a route choice task and their performance demonstrated the benefit of boundaries in real-world state-spaces. This can inform computational theories of planning in machine intelligence by leveraging evidence from human spatial planning experts. Through honing their expert navigation skills over decades, our taxi drivers showed improved performance across the years of experience we sampled (up to 38 years).

